# Developmental Differences in White Matter Microarchitecture in Youth with ADHD: Longitudinal Findings from the ABCD Study

**DOI:** 10.64898/2026.02.20.707054

**Authors:** L. Nate Overholtzer, Katherine L. Bottenhorn, Hedyeh Ahmadi, Sarah L. Karalunas, Bradley S. Peterson, Megan M. Herting

## Abstract

**Background:** Attention-deficit/hyperactivity disorder (ADHD) is the most common neurodevelopmental disorder and is a risk factor for later brain disorders. Here, we characterize the relationship between ADHD status and white matter cellularity across development and examine associations with medication, using a novel biophysical diffusion MRI model in youth aged 9 to 14 years.

Methods:

The ABCD Study® is a longitudinal cohort study with three biennial waves of brain MRI collection. Twenty-seven white matter tracts were delineated using multi-shell diffusion-weighted imaging (DWI) and tractography. Intracellular isotropic (RNI) and directional (RND) diffusion were quantified using the Restriction Spectrum Imaging (RSI) model. Longitudinal linear mixed-effect models characterize the effects of ADHD status and medication use on white matter cellularity across three waves.

Results:

By wave: 9,426 participants at baseline (mean [SD] age: 9.92 [0.63] years; 48.7% Female; 12.2% with ADHD), 6745 participants at 2-year (11.95 [0.65] years; 46.8% Female; 11.3% with ADHD), and 2,483 participants at 4-year (14.07 [0.69] years; 46.0% Female; 11.8% with ADHD). ADHD was associated with decreased RNI in 20 tracts at age 9, with evidence of developmental trajectory differences suggesting attenuation over early adolescence. We found enduring ADHD-associated decreases in RND of 16 tracts spanning ages 9 to 14 years, with methylphenidate effects on 2 tracts. Low-motion sensitivity analyses confirmed robust RNI findings, but not RND findings.

**Conclusions:** ADHD was associated with reductions in isotropic diffusion in white matter tracts, and possibly with complementary reductions in directional diffusion of select tracts. Isotropic diffusion findings suggest atypical glial cellularity in white matter during late childhood.

## Introduction

Attention-deficit hyperactivity disorder (ADHD) is the most common neurodevelopmental disorder, with an estimated population prevalence of 3-10% of children (1–4). ADHD is marked by behavioral issues involving inattention and/or hyperactivity-impulsivity, and which interfere with daily functioning (5). ADHD poses a health burden to pediatric cognitive and social development and is an important indicator for lifetime mental health (6). Despite its high prevalence and significant health burden, neuroimaging has struggled to identify replicable and generalizable brain differences associated with the disorder (7–9), highlighting an opportunity for population-based neuroimaging studies to provide replicable and clinically meaningful insights.

White matter facilitates efficient communication between brain regions, thereby supporting higher-order cognitive processes (e.g., executive functioning, attentional control, and emotion regulation) that are commonly impaired in ADHD (10,11). White matter structure and connectivity have emerged as a central focus among proposed mechanisms underlying ADHD neurocircuitry (12,13). For decades, diffusion tensor imaging (DTI) has been utilized to investigate microstructural differences in white matter. Systematic reviews and meta-analyses have reported ADHD-related differences across the three fiber classes: commissural fibers within the posterior corpus callosum (9,14–16), association fibers including limbic fasciculi and cingulum (9,13,17), and projection pathways involving fronto-striatal circuits and corticospinal tracts (9,13,18). These studies report reduced fractional anisotropy (FA) in individuals with ADHD, a metric often interpreted as “reduced white matter integrity,” thought to reflect delayed or dysregulated myelination.

Notably, DTI oversimplifies the complexities of white matter tissue (19), and advanced biophysical diffusion modeling techniques offer improved histological specificity (20–23). Furthermore, these studies also suffer from other methodological concerns: (A) insufficiently accounting for head motion, which can produce spurious findings and is of particular concern due to ADHD hyperactivity symptoms (15), (B) cross-sectional study designs, which limit inferences about neurodevelopmental trajectories, and (C) a lack of consideration for pharmacologic effects. To date, only two DTI studies have directly examined methylphenidate effects on white matter (24,25). Given the protracted maturation of white matter into the third decade of life (26), longitudinal studies examining the relationship between white matter maturation and ADHD may offer promising targets for intervention.

The primary objective of this study was to characterize the relationship between ADHD and development of white matter microarchitecture across early adolescence using restriction spectrum imaging (RSI), a novel biophysical diffusion model (20,23,27). The secondary objective was to evaluate associations with three major classes of ADHD medications: amphetamine-based, methylphenidate-based, and nonstimulants. Unlike DTI, RSI distinguishes between tissue compartments, providing greater specificity in characterizing diffusion properties reflecting axonal organization versus glial cellularity. To achieve this, we leveraged the landmark Adolescent Brain and Cognitive DevelopmentL Study (ABCD Study®) cohort. Based on DTI literature, we hypothesized that axonal organization of white matter tracts connecting frontal, parietal, and striatal regions would be reduced in youth with ADHD.

## Methods and Materials

### Participants and Procedures

This work uses longitudinal data (Release 5.1; http://dx.doi.org/10.15154/z563-zd24) from the ABCD Study, a United States longitudinal cohort study (28). Participants (N = 11,868) aged 9.00 to 10.99 years old at enrollment were recruited through stratified probability sampling at 21 study sites using school-level rosters or birth registries (29). Details of the conception and recruitment for the ABCD Study have been published by Volkow et al. (2018) and Garavan et al. (2018) (28,29). In brief, enrollment criteria for the ABCD Study required participants to be proficient in English, capable of completing MRI procedures, and to not be affected by any severe medical, sensory, or intellectual impairments (27). The University of California, San Diego, Institutional Review Board (IRB) provided centralized approval for experimental and consent procedures, and each study site obtained local IRB approval. Participants provided written assent, and their legal guardians provided written informed consent.

For this study, we leverage assessments conducted at baseline (2016–2018), 2-year visit (2018–2020), and 4-year visit (2020–2022). The present study’s inclusion/exclusion criteria are visualized with changes to the sample size in Supplemental Figure 1. Participants were excluded if diffusion MRI was not collected at a visit, if T1-weighted or diffusion MRI failed quality control (30), or if incidental radiological findings were present (31). Additional exclusions included missing mental health or medication assessments, incomplete demographic data, and reported current use of antidepressant, antipsychotic, antiepileptic, anxiolytic, or other neurologic or psychiatric medications.

### RSI Acquisition and Processing

A harmonized multi-shell diffusion-weighted imaging protocol was implemented across all study sites, with preprocessing and quality control performed by the ABCD Data Analysis, Informatics, and Resource Center (30). Details of acquisition and processing are provided in *Supplemental Methods*. Average head motion (*mm*) was calculated from framewise displacement (FD) during acquisition (30).

RSI, an advanced biophysical diffusion model, was then applied to quantify diffusion properties of extracellular and intracellular white matter compartments. From this approach, we selected two diffusion metrics within the intracellular compartment: restricted normalized isotropic signal fraction (RNI) and restricted normalized directional signal fraction (RND), both scaled from 0 to 1 (20,21,23). In white matter, RNI measures isotropic diffusion within the restricted (i.e., intracellular) compartment, thought to index glial cell density or size (27), whereas RND reflects anisotropic (directional) diffusion within the restricted (i.e., intracellular) compartment, believed to indicate axonal packing density and myelin integrity (27) (*Figure 1*).

**Figure 1.**
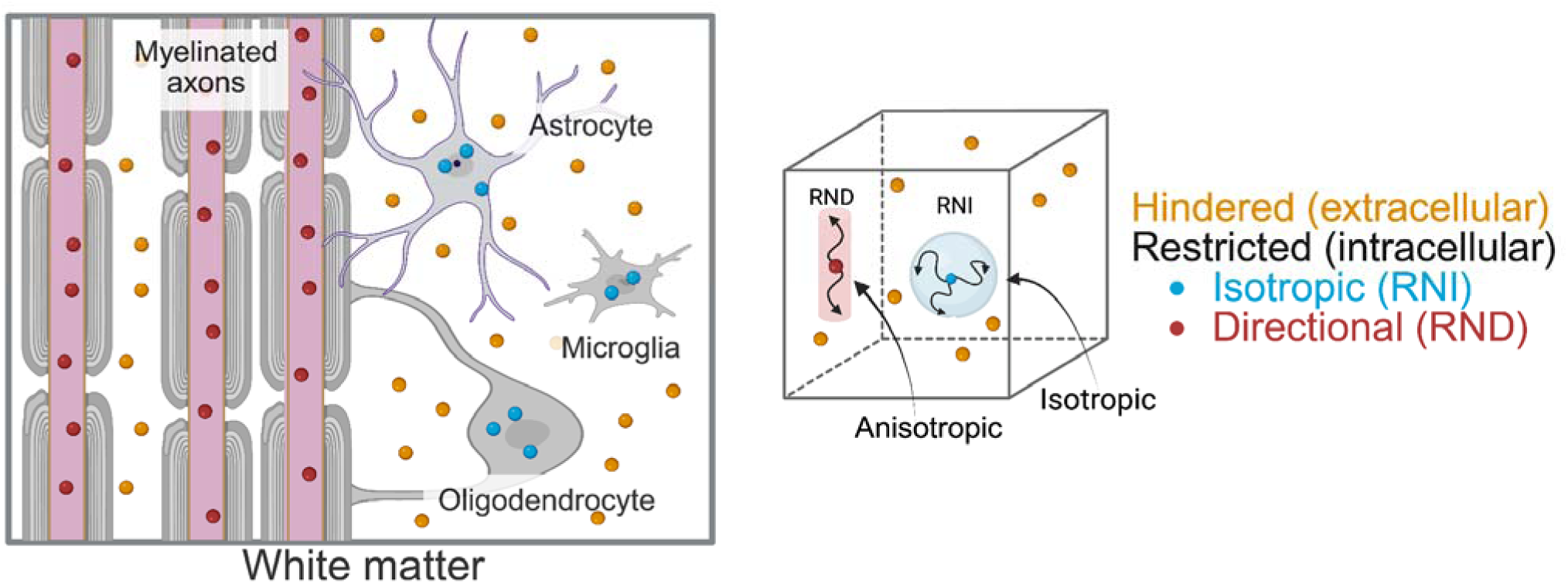
Illustration of the RSI Biophysical Model of Diffusion. Restricted diffusion reflects intracellular water movement. Restricted normalized directional (RND) diffusion indicates anisotropic diffusion along elongated structures, indexing water movement along myelinated axons. Restricted normalized isotropic (RNI) diffusion indicates isotropic diffusion within the cell bodies of support cells, including astrocytes, microglia, and oligodendrocytes.

Twenty-seven white matter tracts of interest were delineated using the probabilistic AtlasTrack Fiber Tract Atlas (32), including three unilateral tracts (corpus callosum, forceps major, and forceps minor) and twelve bilateral tracts (fornix, cingulate cingulum, parahippocampal cingulum, corticospinal [pyramidal] tract, anterior thalamic radiations, uncinate, inferior longitudinal fasciculus, inferior fronto-occipital fasciculus, superior longitudinal fasciculus, superior corticostriate, striatal inferior frontal cortex tract, and inferior frontal–to–superior frontal cortical tract). Mean RNI and RND values computed within each tract mask served as the outcomes in statistical models.

### ADHD Status and Medication Classification

Participants were identified as having ADHD or as non-ADHD controls based on caregiver responses to the computerized *Kiddie Schedule for Affective Disorders and Schizophrenia* (KSADS) and the Medication History Questionnaire (MHQ) (33–35). We used the MHQ to determine participants taking amphetamine-based (AMP), methylphenidate-based (MPH), and nonstimulant (NS) medications, and to exclude individuals taking neurologic or psychiatric medications for conditions other than ADHD. Those meeting ADHD diagnostic criteria and/or reporting ADHD medication use were assigned to the ADHD group for the respective wave. This approach addresses concerns in ABCD that parents or caregivers were asked to report on participant’s current behavior, which could lead to children well-treated by ADHD medications appearing to not meet diagnostic criteria. Some participants exhibited changes in ADHD status across waves, including both onset and remission; *Supplemental Figure 2* illustrates longitudinal participant flow.

### Sociodemographic Data

Sociodemographic covariates included caregiver-reported sex at birth, race/ethnicity, annual household income, and parent education. Time-invariant variables, assessed at baseline, were sex at birth (male, female, intersex male, or intersex female) and race/ethnicity. Caregivers reported participants’ race and ethnicity using a single item for race and a separate item for Hispanic/Latino ethnicity. Participants were classified into one of the following categories: non-Hispanic White (only White), non-Hispanic Black (only Black/African American), Hispanic/Latino (any endorsement of Hispanic/Latino identity), non-Hispanic Asian (Asian Indian, Chinese, Filipino, Japanese, Korean, Vietnamese, or other Asian), or multiracial/other (including American Indian/Native American, Alaska Native, Native Hawaiian, Guamanian, Samoan, other Pacific Islander, other race, or more than one race), or Refuse/Don’t Know (36). Time-varying variables, reassessed at each study wave, included age (in months), a collapsed measure of household income (<$50,000 USD, $50,000–$99,999 USD, ≥$100,000 USD, or Refuse/Don’t Know), and highest household education (less than high school, high school diploma/GED, some college, bachelor’s degree, postgraduate degree, or Refuse/Don’t Know).

### Statistical Analyses

Linear mixed-effects models measured effects of primary predictors for ADHD status and each class of medication use (amphetamine, methylphenidate, and nonstimulant), as well as an ADHD-by-age interaction for group differences in developmental trajectories. Each primary predictor was binary and time-varying, reflecting within-subject changes in medication use and/or ADHD status (*Supplemental Figure 2*). Models were adjusted for sociodemographic covariates and head motion, along with crossed random effects of MRI scanner, family, and subject. Unstandardized β coefficients with 95% confidence intervals, Cohen’s f² effect sizes, and uncorrected and false discovery rate (FDR)–corrected P values for our effects of interest are reported. The variance attributable to random effects, expressed as the intraclass correlation coefficient (ICC), was calculated.

All models were fit with and without the ADHD-by-age interaction term and compared using likelihood ratio tests. Interaction models were preferred for RNI outcomes, and models without the interaction were preferred for RND outcomes. Results from alternative models are in *Supplemental Results*. Age was centered at baseline (9 years) and scaled in years, such that the interaction term reflects the estimated yearly change in diffusion values for the ADHD group relative to the control group. We conducted post hoc estimated marginal means (EMM) analyses to summarize ADHD–control differences in RNI across ages 9.00 through 15.00 years. Sensitivity analyses were also conducted, which excluded participants using a more stringent head motion criteria (FD ≥ 2mm).

## Results

The analytical sample comprised 18,654 observations for 10,526 participants (*Table 1*): 9,426 at baseline (mean [SD] age, 9.92 [0.63] years; 48.7% female; 12.2% with ADHD), 6,745 at 2-year (11.95 [0.65] years; 46.8% female; 11.3% with ADHD), and 2,483 at 4-year (14.07 [0.69] years; 46.0% female; 11.8% with ADHD). In the ADHD group, 658 participants at baseline, 510 at 2-year, and 187 at 4-year reported use of one or more ADHD medications. Full demographic characteristics are presented in *Table 1*, and group-specific characteristics are provided in *Supplemental Table 1*. Across all waves, the ADHD group had more males and race/ethnicity distributions differed; the ADHD group also showed greater head motion at baseline and 2-year. Descriptive trajectories of RNI and RND for each white matter tract are illustrated in *Supplemental Figures 3* and *4*. The variance attributable to random effects is reported in *Supplemental Figure 5*.

**Table 1.**
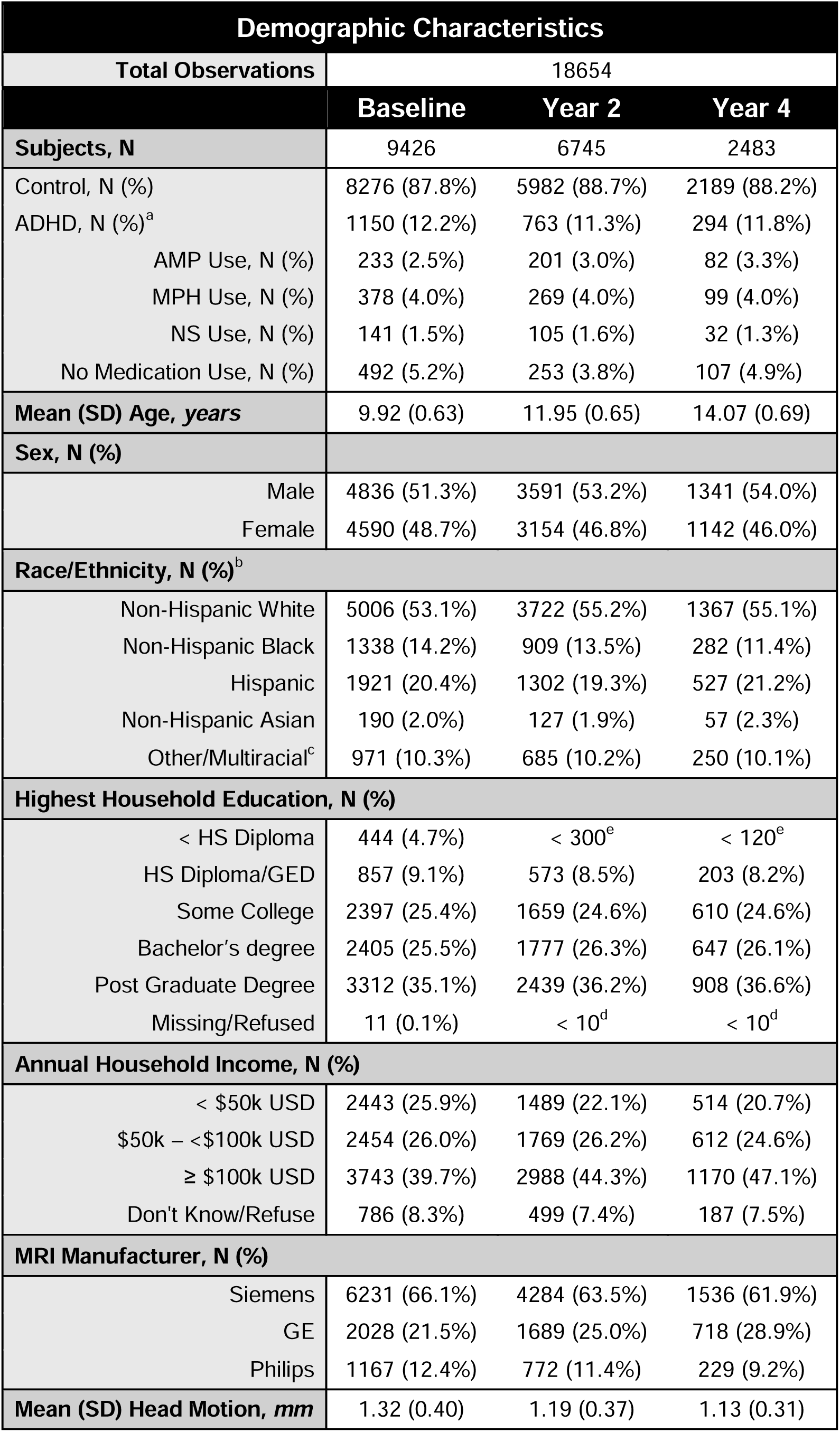
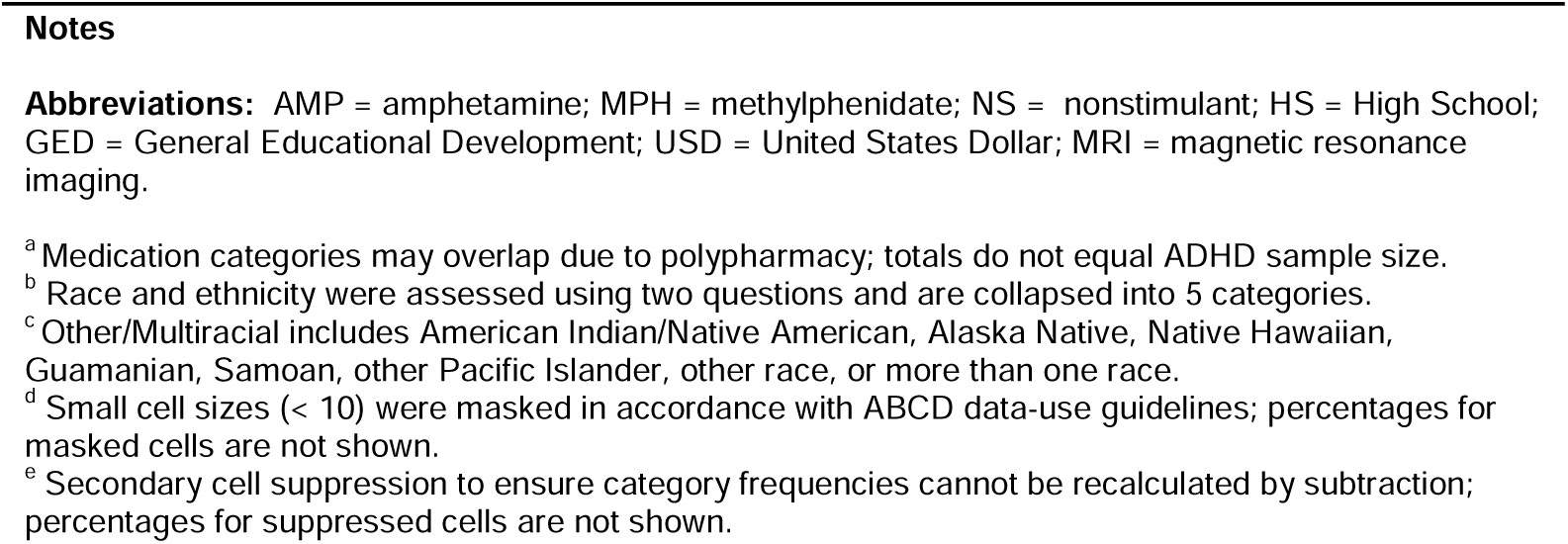
Demographic Characteristics of the Analytical Sample by Wave.

### Intracellular Isotropic Diffusion (RNI)

The ADHD-by-age interaction term improved model fit in 9 of 27 white matter tracts and was retained in all RNI models to facilitate direct comparison. Interactions were observed in the left and right fornix, forceps major, and left corticospinal tract (*P*_FDR_ < .05) (*Figures 2–3*); 5 additional tracts reached uncorrected significance (*P* < .05) (*Figure 2* and *Supplemental Table 2*). All interaction effects were positive, indicating that ADHD-related differences decreased between ages 9 and 14 years; post-hoc estimated marginal means indicate that group differences are non-significant in all tracts by age 13 years (*Supplemental Table 3*).

**Figure 2.**
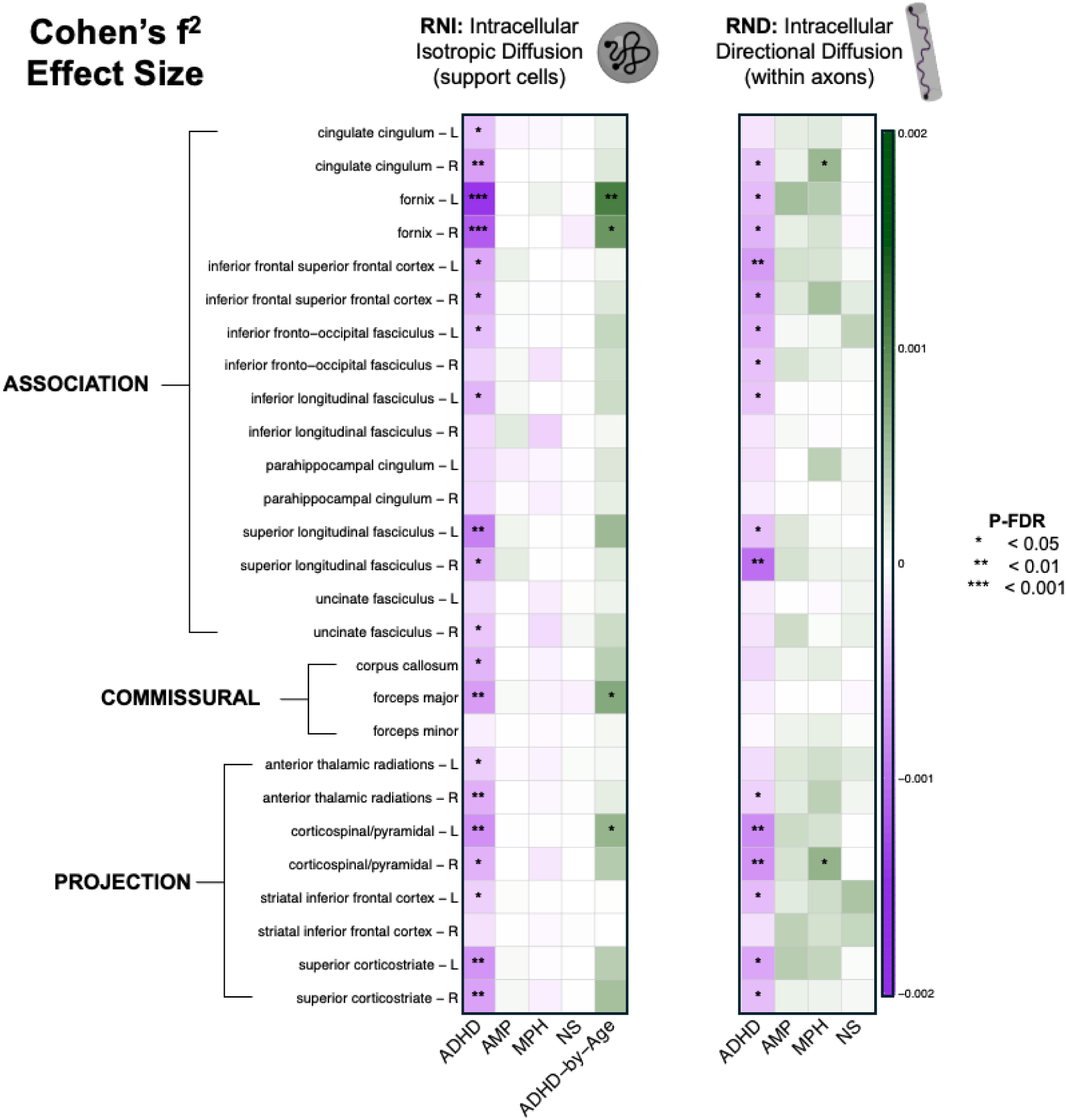
Effect Sizes of ADHD and Medication Associations with White Matter Microarchitecture. Heatmaps display effect sizes (Cohen’s f²) from models examining the associations between ADHD and medication use (AMP, MPH, NS) with RSI measures of white matter microstructure. Restricted normalized isotropic diffusion (RNI) models reflect intracellular diffusion in glial cells of white matter tracts, whereas directional diffusion (RND) models reflect intra-axonal diffusion of tracts. Rows correspond to white matter tracts with left (L) and right (R) hemispheres shown separately. Columns indicate predictors, including ADHD-by-age interaction for RNI. Colors reflect effect direction and magnitude (purple = negative; green = positive). False discovery rate–adjusted significance is denoted: * < .05; ** < .01; *** < .001.

**Figure 3.**
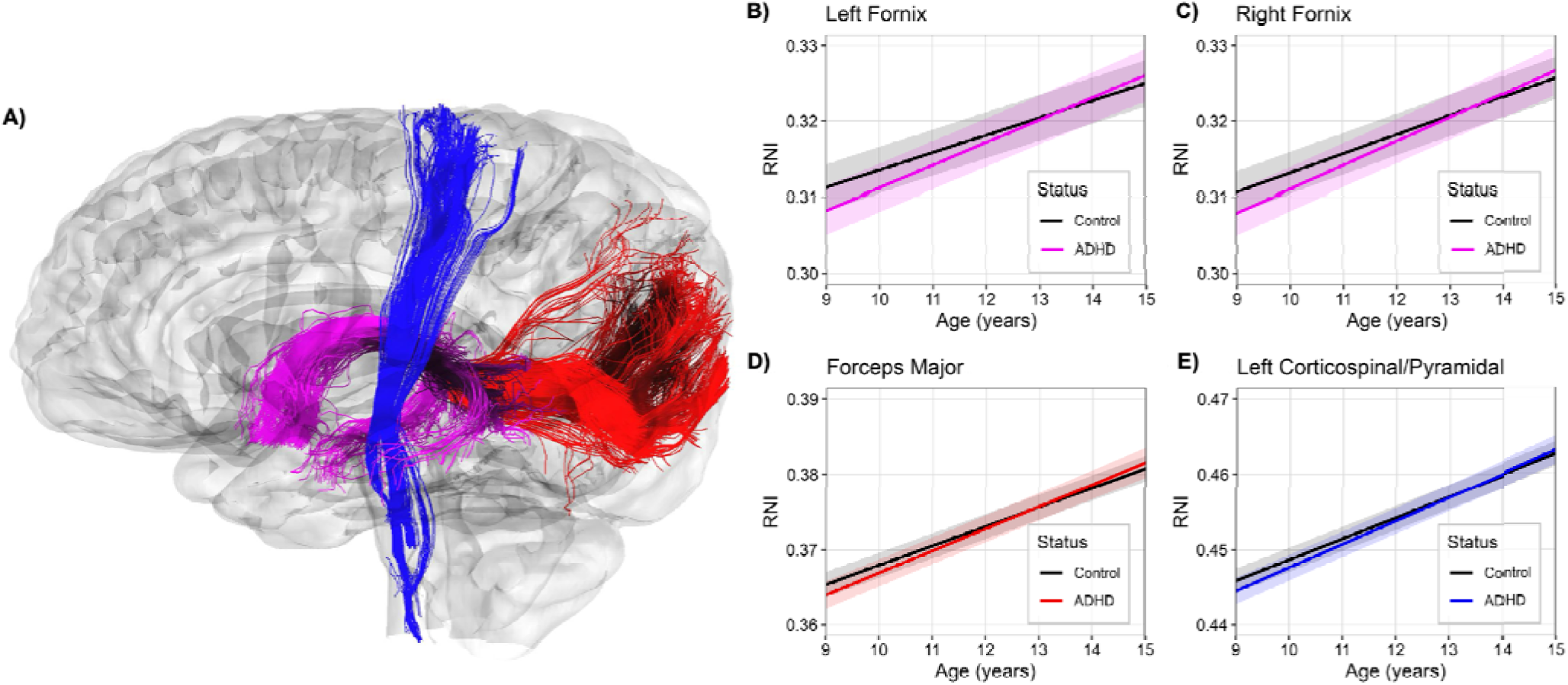
Developmental Trajectory Differences in Isotropic Diffusion Across Four White Matter Tracts. Only four white matter tracts demonstrated a significant ADHD-by-age interaction after FDR-correction, which are visualized in a left lateral view of the brain: left fornix (magenta), right fornix (magenta), forceps major (red), and left corticospinal/pyramidal tract (blue) (**A**). Developmental trajectories of restricted normalized isotropic diffusion (RNI) for each tract are shown for youth with ADHD and controls (**B–D**). Group means and 95% CIs were estimated using linear mixed-effects models. Colored lines indicate the ADHD group, and black lines indicate the control group. Higher RNI values reflect greater intracellular isotropic diffusion, a marker of glial cellularity or morphology. Across all four tracts, youth with ADHD exhibited lower RNI at age 9 but exceed controls by age 14.

Main effects of ADHD on RNI were present in 20 of 27 tracts (*P*_FDR_ < .05), with 4 additional tracts at uncorrected levels (*P* < .05) (*Figure 2* and *Supplemental Table 2*): the left and right parahippocampal cingulum, left uncinate fasciculus, and right inferior fronto-occipital fasciculus. Exceptions (*P* > .05) included the right inferior longitudinal fasciculus, right striatal–inferior frontal cortex, and forceps minor. All ADHD effects were negative, indicating lower tract RNI among youth with ADHD compared with controls at age 9 years. In the 4 tracts with significant interaction terms (left and right fornix, forceps major, and left corticospinal tract), the trajectories of the ADHD and control groups intersected around ages 12.5 to 13.5 years, after which the ADHD group exhibited greater RNI values (*Figure 3*). No corrected associations were significant for medication class, and medication effects were largely nearly-zero for RNI (*Figure 2*). However, methylphenidate displayed 2 uncorrected associations (*P* < .05) in the right inferior longitudinal fasciculus and right uncinate fasciculus (*Supplemental Table 2*).

### Intracellular Directional Diffusion (RND)

For RND models, the parsimonious model excluding the ADHD-by-age interaction was preferred, as inclusion improved model fit in only 1 tract. Main effects of ADHD on RND were observed in 16 of 27 tracts (*P*_FDR_ < .05), indicating lower RND among youth with ADHD compared to controls across ages 9 through 14 years (*Figure 2* and *Supplemental Table 4*). Affected tracts included the right cingulate cingulum, bilateral fornix, bilateral inferior frontal–superior frontal cortex, left inferior fronto-occipital fasciculus, left inferior longitudinal fasciculus, bilateral superior longitudinal fasciculus, right anterior thalamic radiation, bilateral corticospinal tracts, left striatal–inferior frontal cortex, and bilateral superior corticostriate tracts.

Methylphenidate use was associated with higher RND in 2 tracts (*P*_FDR_ < .05): the right cingulate cingulum and right corticospinal tract (*Figure 2* and *Supplemental Table 4*). Additional uncorrected associations (*P* < .05) were observed in 6 tracts for amphetamine, 8 for methylphenidate, and 3 for nonstimulants (*Supplemental Table 4*). All corrected and uncorrected associations on RND were positive, indicating attenuation of the ADHD phenotype.

### Sensitivity Analysis

After excluding participants with FD ≥ 2mm, the sensitivity analysis sample included 8,939 at baseline, 6,517 at 2-year, and 2,438 at 4-year (*Supplemental Table 5*). The prevalence of ADHD in the sensitivity sample (11.9% at baseline, 11.1% at 2-year, and 11.9% at 4-year) was similar to the primary analytical sample (differences ≤0.3%), but with slightly greater attrition in the ADHD group. Distributions of medication use, sociodemographic characteristics, and MRI manufacturer were comparable to those of the full sample across all time points (*Supplemental Table 5*).

After excluding high-motion participants, the ADHD-by-age interaction in left and right fornix RNI remained (*P*_FDR_ < .05) (*Supplemental Figure 6* and *Supplemental Table 6*). Main effects of ADHD on RNI persisted in 13 tracts: bilateral fornix, bilateral cingulate cingulum, bilateral corticospinal tracts, left inferior longitudinal fasciculus, forceps major, corpus callosum, bilateral superior longitudinal fasciculus, and bilateral superior corticostriate tracts (*P*_FDR_ < .05).

In the RND sensitivity analysis, no associations survived FDR correction (*P*_FDR_ ≥ .05; *Supplemental Figure 7* and *Supplemental Table 7*). However, uncorrected associations (*P* < .05) were observed for ADHD in 10 tracts, as well as multiple uncorrected associations for amphetamine, methylphenidate, and nonstimulant use.

## Discussion

This study is the first to demonstrate that biophysical modeling of diffusion using RSI may enhance our understanding of compartment-specific white matter features associated with ADHD. Decreased isotropic diffusion (i.e., glial cellularity), more so than directional diffusion (i.e., axonal organization/myelin), differentiated white matter microarchitecture in late childhood but not mid-adolescence—highlighting a transient developmental window. We found robust evidence of widespread ADHD-related reductions in intracellular isotropic diffusion at age 9 years, with reductions in 20 tracts in the primary analyses and persisting in 13 tracts after stringent motion analyses. Additionally, group trajectory differences suggest that these late childhood deficits attenuate or reverse in early adolescence for those with ADHD. Decreases in directional diffusion associated with ADHD were enduring in 16 tracts across ages 9 through 14 years, which may indicate that myelin caliber is another component of ADHD pathophysiology. However, these findings were not robust in stringent motion analyses.

Isotropic diffusion within the restricted compartment reflects uniform, directionally independent intracellular water movement, which is thought to index changes in the number or morphology of glial cells (i.e., microglia, astrocytes, and oligodendrocytes) (20,27). Reduced isotropic diffusion may indicate fewer or less mature glia in those with ADHD during late childhood. This aligns with emerging evidence for a role of glial dysfunction in ADHD (37,38), including microglia dyshomeostasis (39,40), astrocyte regulation of neurotransmission and synapse formation (41–44), and oligodendrocyte hypomyelination (12,38). Glial cells play roles beyond mere support, actively contributing to brain development through early-life circuit formation, pediatric synaptic patterning and pruning, and lifetime refinement of connectivity (37). The ST3GAL3 gene—a regulator of oligodendrocyte-mediated myelination—has been identified in GWAS analyses of ADHD (45). Rodent knock out models lead to reduced proliferation of oligodendrocyte precursors and decreased hippocampal arborization (38,46), which may also contextualize our fornix findings.

Group developmental trajectories of isotropic diffusion suggest that late childhood differences in youth with ADHD diminish or reverse during early adolescence in impaired tracts. A small cohort has reported similar longitudinal trajectories of more rapid development in FA (47). Importantly, our work suggests that childhood deficits attenuate during early adolescence, and a highly powered sample may be necessary to detect such developmental heterogeneity that could be obscured in mixed-age pediatric cohorts. For the bilateral fornix, forceps major, and left corticospinal tract, trajectories converge around ages 12.5 to 13.5 years, which may support a neurodevelopmental ‘catch-up’ period for the ADHD group (6,48). Alternatively, the rapid gains in isotropic diffusion by the ADHD group during early adolescence may reflect changes that index neuroinflammation/gliosis (27,37,38), rather than simply ‘catch-up.’

The strongest ADHD-related effects were observed in isotropic diffusion of the bilateral fornix. The expanded limbic system framework builds beyond the Papez circuit to include of cortico–limbo–thalamo–cortical circuits connected by multiple major white matter tracts (49). Two principal pathways of the fornix exist: (A) precommissural fibers project to the septal nuclei and basal forebrain, influencing mesolimbic regions; and (B) postcommissural fibers project to hypothalamic and thalamic nuclei (49). While few studies have implicated the fornix, these findings are supported by recent ABCD work proposing a role of limbic dysregulation in ADHD (50,51). Such dysfunction may manifest in emotional dysregulation, a common and clinically impairing feature of ADHD (52,53). However, emotional dysregulation symptoms in ADHD may also exist within a broader transdiagnostic symptom domain (54,55), and findings within the fornix may be attributable to general transdiagnostic psychopathology, rather than ADHD-specific (56).

Intracellular directional (anisotropic) diffusion reflects intra-axonal water movement along elongated cylindrical structures and is believed to index axonal organization and/or myelination (20,27). In this study, ADHD was associated with possible directional diffusion reductions in select association and projection pathways spanning ages 9 through 14 years, nearly all of which also demonstrated isotropic diffusion effects. In contrast to our isotropic findings, medication use, specifically stimulant classes, was as an important contributor to the explained variance in directional diffusion of tracts. Medication effects were broadly positive, offsetting ADHD-related decreases; only methylphenidate displayed two significant associations in the right cingulate cingulum and right corticospinal tract. Yet, none of the observed effects on directional diffusion passed FDR correction in the stringent low-motion analyses. These discrepancies may reflect: (A) motion influencing spurious primary results, (B) retention bias, as unmeasured ADHD heterogeneity (e.g., severity, hyperactivity, poor medication response) could relate to meeting the low-motion thresholds, or (C) the stringency of FDR correction, which fails to account for non-independence amongst tracts. Accordingly, any findings from primary analyses related to directional diffusion should be interpreted cautiously.

### Strengths and Limitations

Our study capitalizes on the population-based ABCD Study cohort to examine longitudinal associations between ADHD and white matter microarchitecture across six years of childhood and adolescence, although causal inferences should not be made. Prior ABCD Study analyses using DTI have reported ADHD to be associated with reduced FA in frontoparietal, fronto-thalamic, and fronto-subcortical pathways (17,51,57). In comparison, our longitudinal findings suggest that RSI offers novel insights into in white matter microarchitecture development in those affected by ADHD. Moreover, few prior diffusion studies have accounted for medication use, which we demonstrate to be an important source of variance in directional diffusion. We also applied stringent quality-control criteria, including sensitivity analyses in a low-motion subsample.

A primary limitation is that analyses are restricted to the first few ABCD Study imaging waves. Most participants did not yet have third-wave MRI data, and collection of second-wave MRI data was impacted by COVID-19 pandemic. Future waves will be essential for characterizing developmental change, particularly trajectories of isotropic diffusion. Another limitation is KSADS diagnoses rely on parent report rather than clinician-administered interviews, although prior research supports their validity and reliability (58). While many children undoubtedly have ADHD, some children with attention and impulse control problems related to other presentations may also be captured by KSADS parent-report. Our observed effect sizes were very small. Nevertheless, small effects can yield meaningful population-level consequences (59), and developmental neuroimaging studies show reduced effect sizes due to variance absorbed by random effects (e.g., scanner, family, and subject).

Future research should include measures of ADHD heterogeneity, which may contribute to differences in our sensitivity analyses. Additionally, a small body of DTI literature suggests sex may influence ADHD-related differences in white matter microstructure (60–62); exploring the modifying effects of sex will be an important future direction. Finally, we cautiously hypothesize that early alterations in isotropic diffusion, possibly reflecting the coordinated role of glia in white matter plasticity during development, may set the stage for the subsequent adolescent differences in directional diffusion, which indexes axonal organization and coherence of white matter pathways. Future work should evaluate this with models suited for causal and temporal inference.

### Conclusions

This cohort study found robust evidence that ADHD was associated with decreased isotropic diffusion across association, commissural, and projection white matter tracts in late childhood, consistent with an altered glial environment; these differences attenuated or reversed during early adolescence. We also observed complementary, enduring ADHD-related decreases in directional diffusion in select tracts spanning ages of 9 to 14 years, consistent with altered myelin caliber; however, these findings were not robust in low-motion analyses. Altogether, we highlight the potential of RSI in advancing our understanding of white matter microarchitecture in ADHD and underscore early adolescence as a critical developmental window for elucidating cellular mechanisms and informing interventions.

## Acknowledgments

A special thanks to the participants and families of the ABCD Study. We would like to acknowledge the ABCD Consortium staff for their efforts in collecting data. We would also like to acknowledge Alethea de Jesus, who contributed to the data cleaning and formatting of the ABCD tabulated data prior to data analysis.

## Author Contributions

L. Nate Overholtzer: Conceptualization, Data Curation, Formal Analysis, Investigation, Methodology, Visualization, Writing – Original Draft Preparation, Writing – Review & Editing.

Katherine L. Bottenhorn: Writing – Review & Editing.

Sarah L. Karalunas: Conceptualization, Writing – Review & Editing.

Bradley S. Peterson: Conceptualization, Writing – Review & Editing.

Hedyeh Ahmadi: Methodology, Validation, Writing – Review & Editing.

Megan M. Herting: Funding acquisition, Conceptualization, Methodology, Supervision, Writing – Review & Editing.

## Data Availability Statement

Data used in the preparation of this article were obtained from the Adolescent Brain Cognitive Development (ABCD) Study (https://abcdstudy.org), held in the NIMH Data Archive (NDA). This is a multisite, longitudinal study designed to recruit more than 10,000 children aged 9–10 and follow them over 10 years into early adulthood. The ABCD Study is supported by the National Institutes of Health Grants [U01DA041022, U01DA041028, U01DA041048, U01DA041089, U01DA041106, U01DA041117, U01DA041120, U01DA041134, U01DA041148, U01DA041156, U01DA041174, U24DA041123, U24DA041147]. A complete list of supporters is available at https://abcdstudy.org/nih-collaborators. A list of participating sites and a comprehensive list of study investigators can be found at https://abcdstudy.org/principal-investigators.html. ABCD consortium investigators designed and implemented the study and/or provided data, but did not necessarily participate in the analysis or writing of this report. This manuscript reflects the views of the authors and may not reflect the opinions or views of the NIH or ABCD consortium investigators. The ABCD data repository grows and changes over time. Qualified researchers can request access to ABCD shared data from the ABCD Data Access Committee (DAC). L. Nate Overholtzer had full access to all the data in the study and takes responsibility for the integrity and accuracy of the data analysis.

