## Supplemental Text and Figures for "Developmental Differences in White Matter Microarchitecture in Youth with ADHD: Longitudinal Findings from the ABCD Study"

This document includes:

- Supplemental Methods
- Supplemental Results
- Supplemental References
- Supplemental Figure 1: Participant Selection Diagram
- Supplemental Figure 2: Flow of Participants Across Study Waves
- Supplemental Figure 3: Average Developmental Trajectories of RNI by Tract
- Supplemental Figure 4: Average Developmental Trajectories of RND by Tract
- Supplemental Figure 5: Random Effects in RNI and RND Models Across White Matter Tracts
- Supplemental Figure 6: Comparison of RNI Model Results Between Primary and Sensitivity Analyses
- Supplemental Figure 7: Comparison of RND Model Results Between Primary and Sensitivity Analyses

**Supplemental Methods**

**Participants and Procedures**

We leverage assessments conducted at baseline (2016-2018), 2-year visit (2018-2020), and 4-year visits (November 2020-January 2022). Data collection for the 2-year visit overlapped with the COVID-19 pandemic, during which some participants underwent modified protocols that did not allow for completion of imaging or other assessments. At the time of the ABCD 5.1 data release, 4-year data collection was incomplete, with most participants’ assessments not yet available.

**Restricted Spectrum Imaging (RSI) Acquisition and Processing**

A harmonized scanning protocol was used across study sites, employing either Siemens, Philips, or GE 3T MRI scanners; details are published in Casey et al. (2018) (1). DWI was acquired in the axial plane at a voxel resolution of 1.7 mm³ with a multiband acceleration factor of 3. The ABCD multi-shell diffusion acquisition protocol includes seven b=0 images and ninety-six gradient directions at four non-zero b-values (6 with b = 500 s/mm², 15 with b = 1000 s/mm², 15 with b = 2000 s/mm², and 60 with b = 3000 s/mm²). Preprocessing steps include: 1) eddy current and motion correction, 2) B0-distortion correction, 3) field inhomogeneity correction, 4) resampling to 1.7 mm³ resolution, 5) alignment to atlas space, and 6) manual and automated quality control; details are provided in Hagler et al. (2019) (2). For our analyses, we used data from participants that the ABCD Data Analysis, Informatics, and Resource Center rated as acceptable image quality (*imgincl_t1w_include* = 1 and *imgincl_dmri_include* = 1) and free of abnormal clinical findings (*mrif_score* = 1 or 2). The ABCD Study collects relatively few b=1000 diffusion-weighted volumes, which is an important consideration since DTI relies heavily on this shell; whereas, RSI leverages the full multi-shell acquisition in its model.

**Sociodemographic Data**

Age was calculated from a deliberately masked birthdate (±15 days of uncertainty). For 19 participants with missing income or parent education data at 2-year or 4-year, the most recently reported value was carried forward. To address small cell sizes in statistical modeling, categories were combined as follows: non-Hispanic Asian and multiracial/other were merged, and "Don’t Know/Refuse to Answer" was grouped by household income.

**Statistical Analysis**

**Linear mixed-effect models**

Linear mixed-effects models were conducted using R, version 4.4.1, with the *lme4* package, version 1.1-35.5 (3). Model diagnostics were evaluated using the *performance* package, version 0.12.2, to confirm that the models met the assumptions of linearity, normality, homoscedasticity, independence of residuals, and validity of the random effects structure (4). Intraclass correlation coefficients were calculated to quantify the proportion of variance attributable to each grouping factor (scanner, family, and subject).

Primary predictors included ADHD status (ADHD vs non-ADHD Control), amphetamine use (yes vs no), methylphenidate use (yes vs no), nonstimulant use (yes vs no), and an ADHD-by-age interaction term to model differential developmental trajectories by group status. Outcomes were isotropic (restricted normalized isotropic [RNI]) and directional (restricted normalized directional [RND]) diffusion metrics derived from the Restriction Spectrum Imaging model, computed across the 27 white matter tracts. Random intercepts (i.e., crossed random effects) were included for MRI scanner serial number, family ID, and subject ID to account for scanner, family relatedness, and repeated measures of subjects.

Model selection followed a stepwise approach. All models were initially fit with an ADHD-by-age interaction term and then refit, excluding the interaction, using maximum likelihood estimation. For each outcome, models were compared using likelihood ratio tests. For RNI outcomes, the interaction model was selected; for RND outcomes, the more parsimonious model without the interaction was selected. For the sake of completeness but not interpretation, results from alternative models are reported in *Supplemental Results* as well as in *Supplemental Tables 8* and *9*.

***Statistical Model Equations for Preferred Model:***

Eq 1. RNI:

White Matter Tract RNI ~ ADHD_Yes/No_ + Age + Age-by-ADHD_Yes/No_ + AMP_Yes/No_ + MPH_Yes/No_ + NS_Yes/No_ + Sex + Race/Ethnicity + Household Income + Parent Education + MRI Manufacturer + (1|MRI Serial No.) + (1|Family ID) + (1|Subject ID)

Eq 2. RND:

White Matter Tract RND ~ ADHD_Yes/No_ + Age + AMP_Yes/No_ + MPH_Yes/No_ + NS_Yes/No_ + Sex + Race/Ethnicity + Household Income + Parent Education + MRI Manufacturer + (1|MRI Serial No.) + (1|Family ID) + (1|Subject ID)

We report unstandardized beta coefficients with 95% confidence intervals calculated using the Wald method, which represent the estimated increase or decrease in RNI or RND units (scaled 0 to 1, normalized in the RSI model), as well as Cohen’s f2. Although Cohen’s f² is normally reported without directionality, we display it with sign to convey whether the effect was positive or negative, while retaining the magnitude as the measure of effect size. False discovery rate (FDR) correction was applied separately for each RSI metric (RNI and RND) and each predictor of interest (ADHD, AMP, MPH, NS, ADHD-by-age). Within each combination, P-values (*P*) were corrected across the 27 white matter tracts using the Benjamini-Hochberg procedure (5), such that each p-value was adjusted against 26 other tracts for the respective predictor. An FDR-corrected P-value (*P*_FDR_) less than 0.05 was used as the threshold of significance.

**Post Hoc Testing of ADHD-by-Age Interaction**

The *emmeans* package was used to obtained group-level estimated marginal means (EMMs) and the ADHD–Control contrast from linear mixed-effect models (6), while adjusting for fixed-effect confounders as proportionally-weighted nuisance variables. The EMMs and group contrast were computed for ages 9.00 through 15.00 years, with inference testing based on asymptotic Wald tests. Negative contrast values indicate lower RNI in the ADHD group relative to the control group.

**Sensitivity Analyses**

Although framewise displacement (FD), an index of average head motion, was included as a covariate in our primary analyses, additional sensitivity analyses were conducted to further assess the robustness of findings. Head motion is known to be associated with both ADHD diagnosis and diffusion MRI (dMRI) signal quality (7), and may act as either a confounder or a mediator of observed effects. We applied more stringent inclusion criteria, excluding participants with an average FD ≥ 2.0 mm during dMRI acquisition, and repeated the analyses in this subsample using the preferred model type. This threshold is more conservative than the standard criteria used by the ABCD Data Analysis and Informatics Resource Center (DAIRC). Sensitivity results are presented in the main manuscript, with corresponding figures included in *Supplemental Figures 6* and *7*.

In some cases, DWI acquisitions were interrupted and restarted midway through the scan. Although the consortium used the FSL *topup* command to correct B0 distortion fields, these interruptions may inflate FD estimates without reflecting true motion.

**Supplemental Results**

These results provide context from the nonpreferred (i.e., alternative) modelling approaches and are presented for completeness. The authors recommend interpreting the following analyses as exploratory and in comparison, to the preferred modelling approach.

Results from the nonpreferred RNI models, which excluded the ADHD-by-age interaction term, are presented in *Supplemental Table 8*. In brief, ADHD was associated with significantly lower RNI values in 9 white matter tracts (*P*_FDR_ < .05).

Results from the nonpreferred RND models, with an ADHD-by-age interaction term, are presented in *Supplemental Table 9*. In brief, there were significant effects of ADHD on RND in 6 white matter tracts and of methylphenidate on RND in 3 white matter tracts (*P*_FDR_ < .05). No significant ADHD-by-age interaction effects were observed, and the directionality of these effects varied by tract, with no consistent pattern emerging.

**Supplemental Figure 1.** **Participant Selection Diagram.** This flow diagram depicts analytical and sensitivity sample selection from the ABCD Study Population (N = 11,868) by study wave at baseline, 2-year, and 4-year follow-up visits. Exclusions were applied sequentially for: (A) missing diffusion MRI data, (B) unusable structural or diffusion scans, incidental findings, (C) missing mental health, medication, or demographic information, and (D) non-ADHD neuropsychiatric medication. The final analytic sample was further restricted in sensitivity analyses to youth with average framewise displacement < 2 mm.

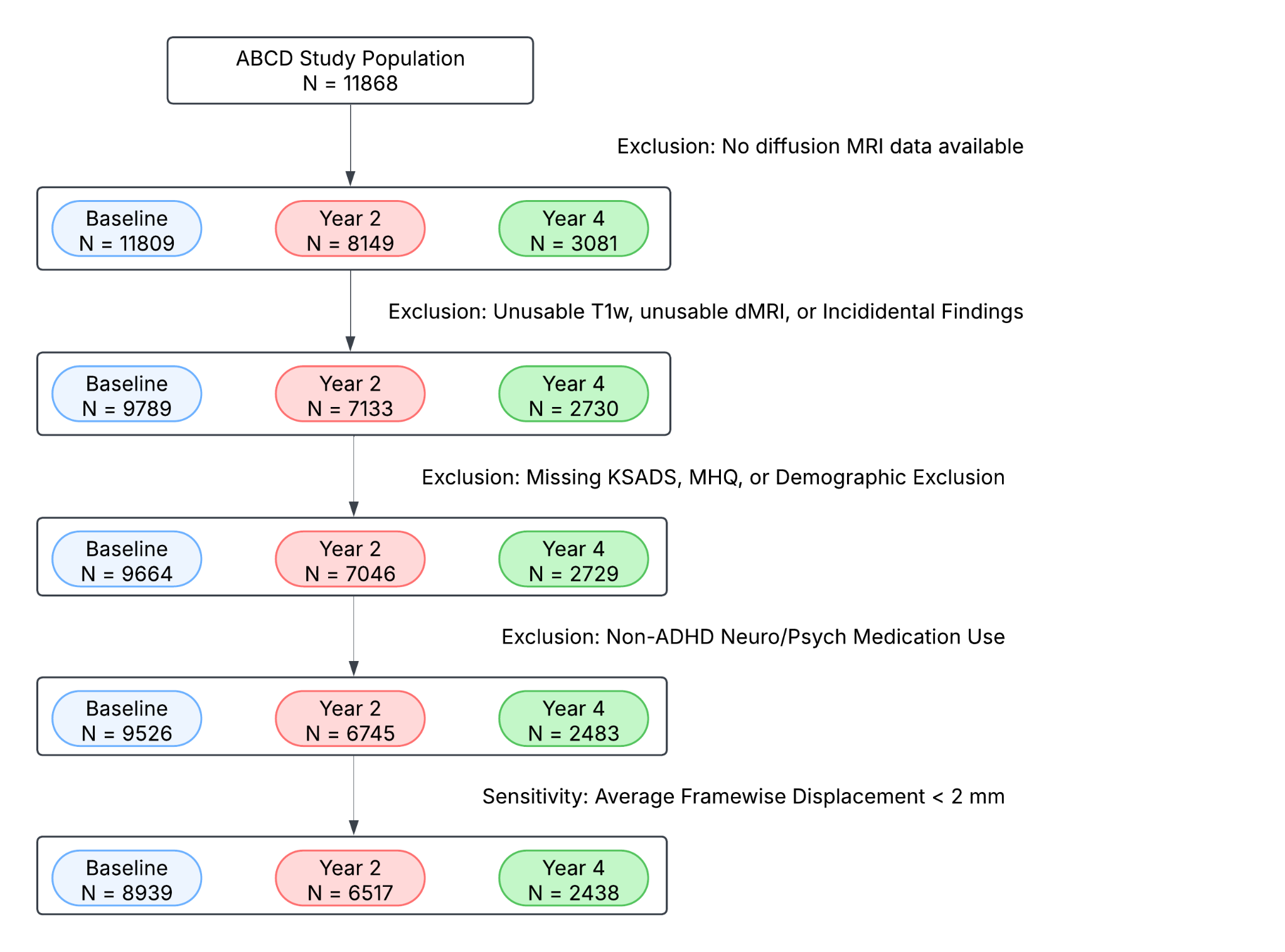

**Supplemental Figure 2.** **Flow of Participants Across Study Visits.** Flow of participants with ADHD and controls at baseline (ages 9–10 years), 2-year follow-up (ages 11–12 years), and 4-year follow-up (ages 13–14 years), including those excluded, with missing data, or missing study waves. Data collection for the 2-year visit coincided with the COVID-19 pandemic, and data collection for the 4-year visit was ongoing at the time of ABCD Data Release 5.1.

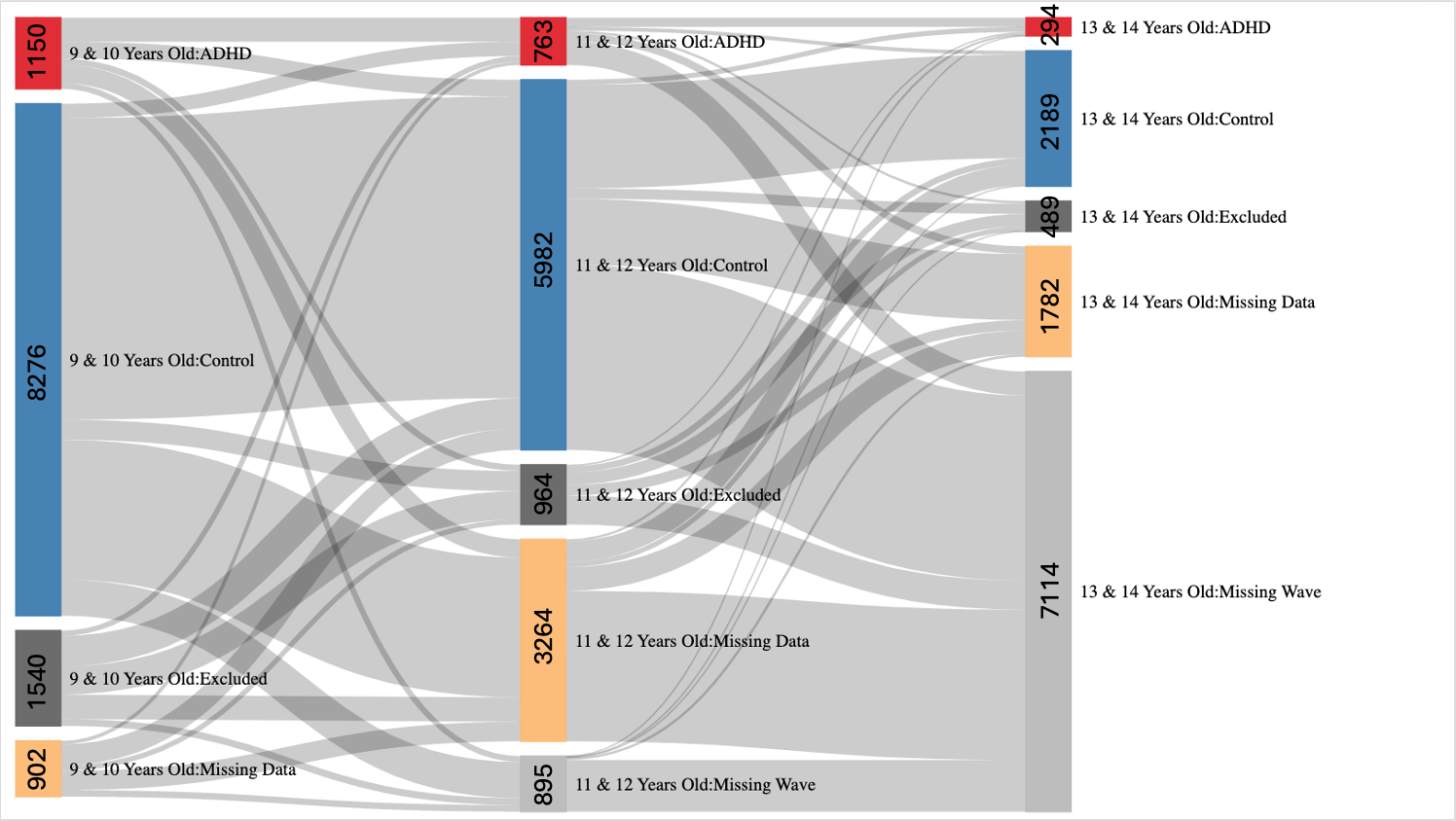

**Supplemental Figure 3.** **Average Developmental Trajectories of RNI by Tract.** General additive models were used to visualize the trajectory of RND development for each tract across ages 9 through 14 years for control youth (red) and youth with ADHD (blue).

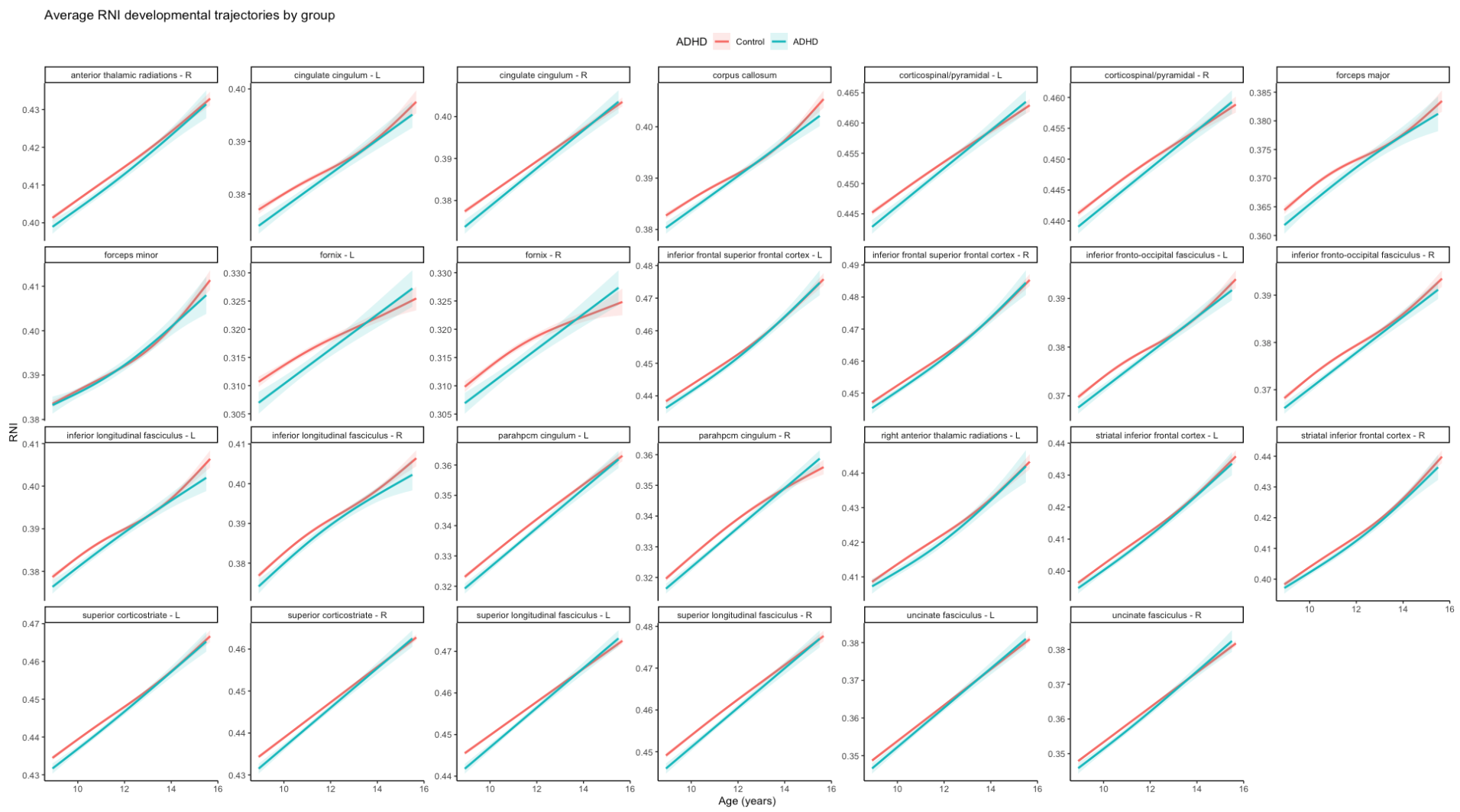

**Supplemental Figure 4.** **Average Developmental Trajectories of RND by Tract.** General additive models were used to visualize the trajectory of RNI development for each tract across ages 9 through 14 years for control youth (red) and youth with ADHD (blue).

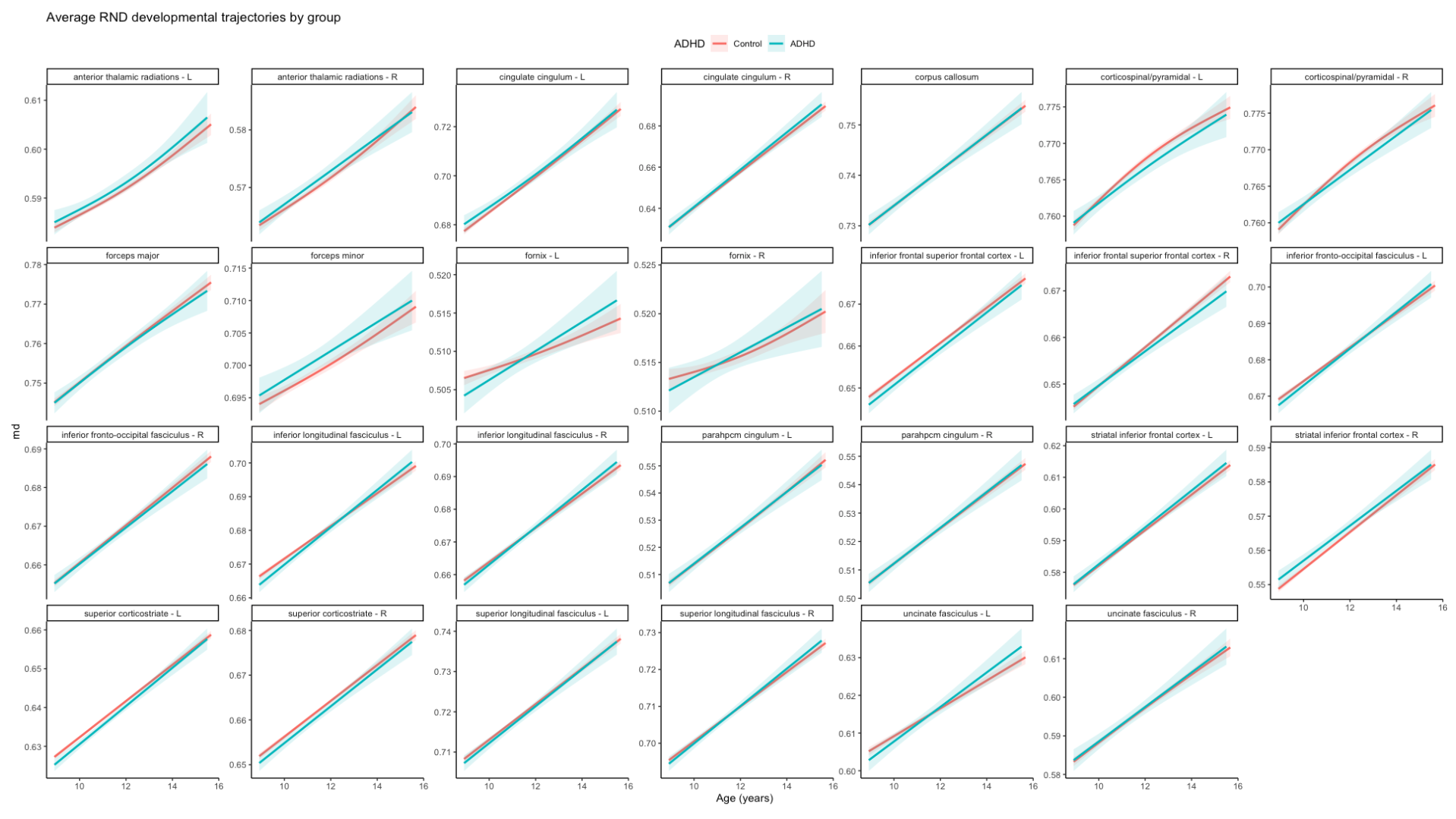

**Supplemental Figure 5. Random Effects in RNI and RND Models Across White Matter Tracts.** Stacked bar plots display the intraclass correlation coefficient (ICC) decomposed by specific random effects of subject (green), family (orange), and scanner (blue) for RNI models by tract (**Top**) and RND models by tract (**Bottom**). Tracts with lower ICC values and smaller subject-level contributions reflect greater developmental variability.

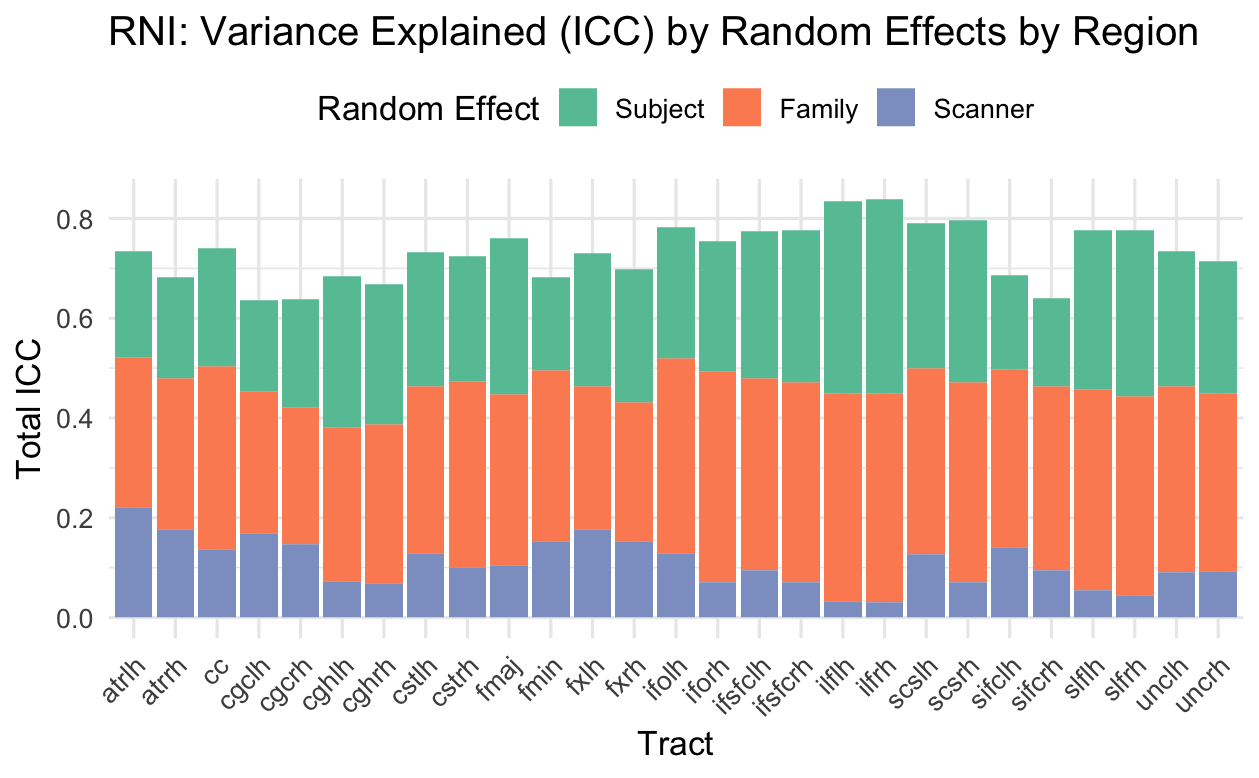

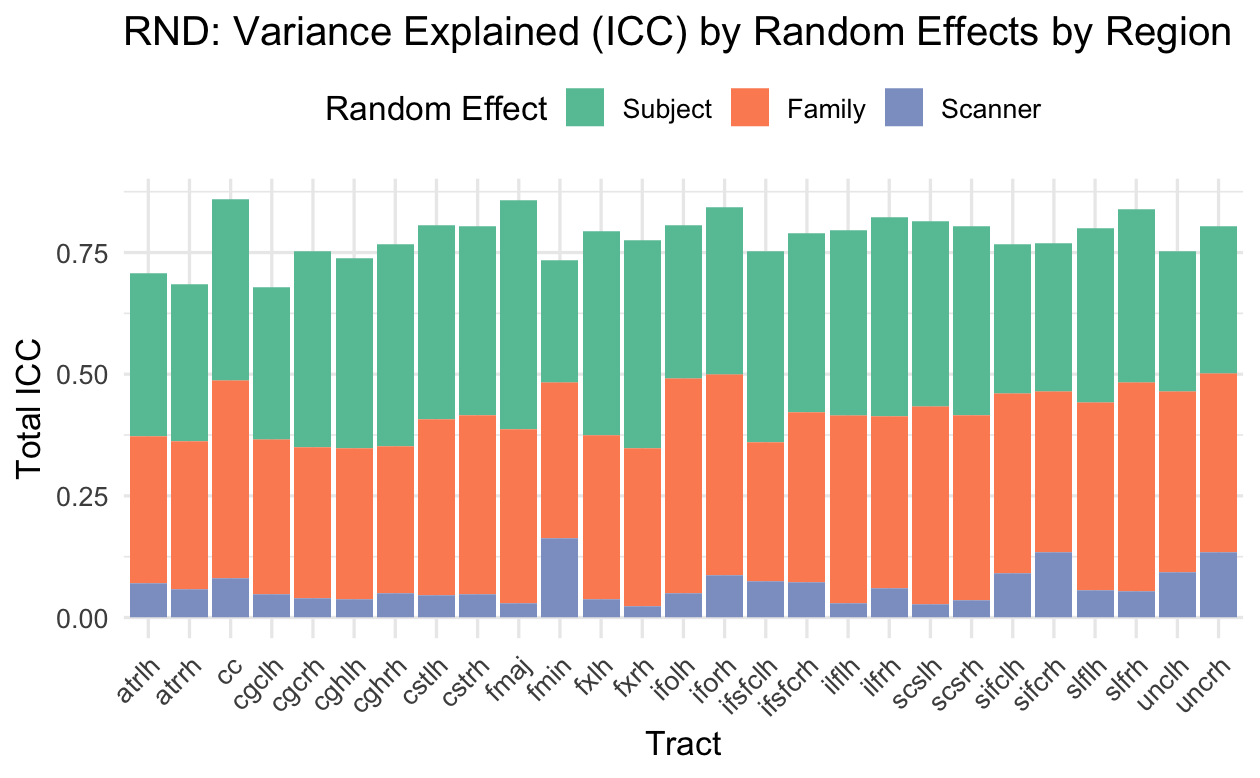

**Supplemental Figure 6. Comparison of RNI Model Results Between Primary and Sensitivity Analyses.** Heatmaps display effect sizes (Cohen’s f²) comparing results from RNI models in the Primary Analysis (Observations = 18654) and Sensitivity Analysis (Observations = 17894). Rows correspond to the RNI of white matter tracts with left (L) and right (R) hemispheres shown separately. Columns indicate predictors, including ADHD, AMP, MPH, NS, and an ADHD-by-age interaction. Colors reflect effect direction and magnitude (purple = negative; green = positive). False discovery rate–adjusted significance is denoted: * < .05; ** < .01; *** < .001.

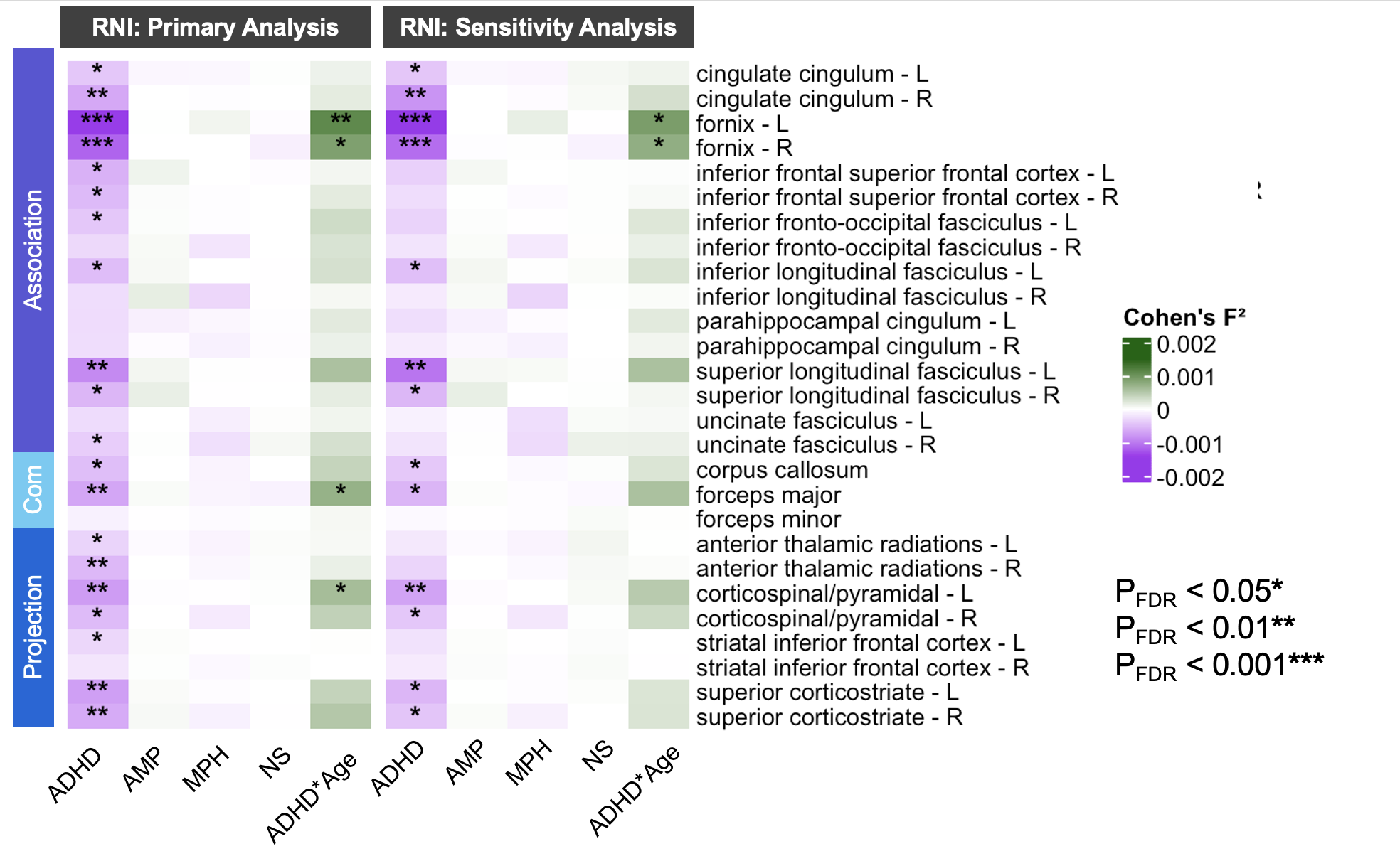

**Supplemental Figure 7: Comparison of RND Model Results Between Primary and Sensitivity Analyses.**

Heatmaps display effect sizes (Cohen’s f²) comparing results from RND models in the Primary Analysis (Observations = 18654) and Sensitivity Analysis (Observations = 17894). Rows correspond to RND of white matter tracts with left (L) and right (R) hemispheres shown separately. Columns indicate predictors, including ADHD, AMP, MPH, and NS. Colors reflect effect direction and magnitude (purple = negative; green = positive). False discovery rate–adjusted significance is denoted: * < .05; ** < .01; *** < .001.

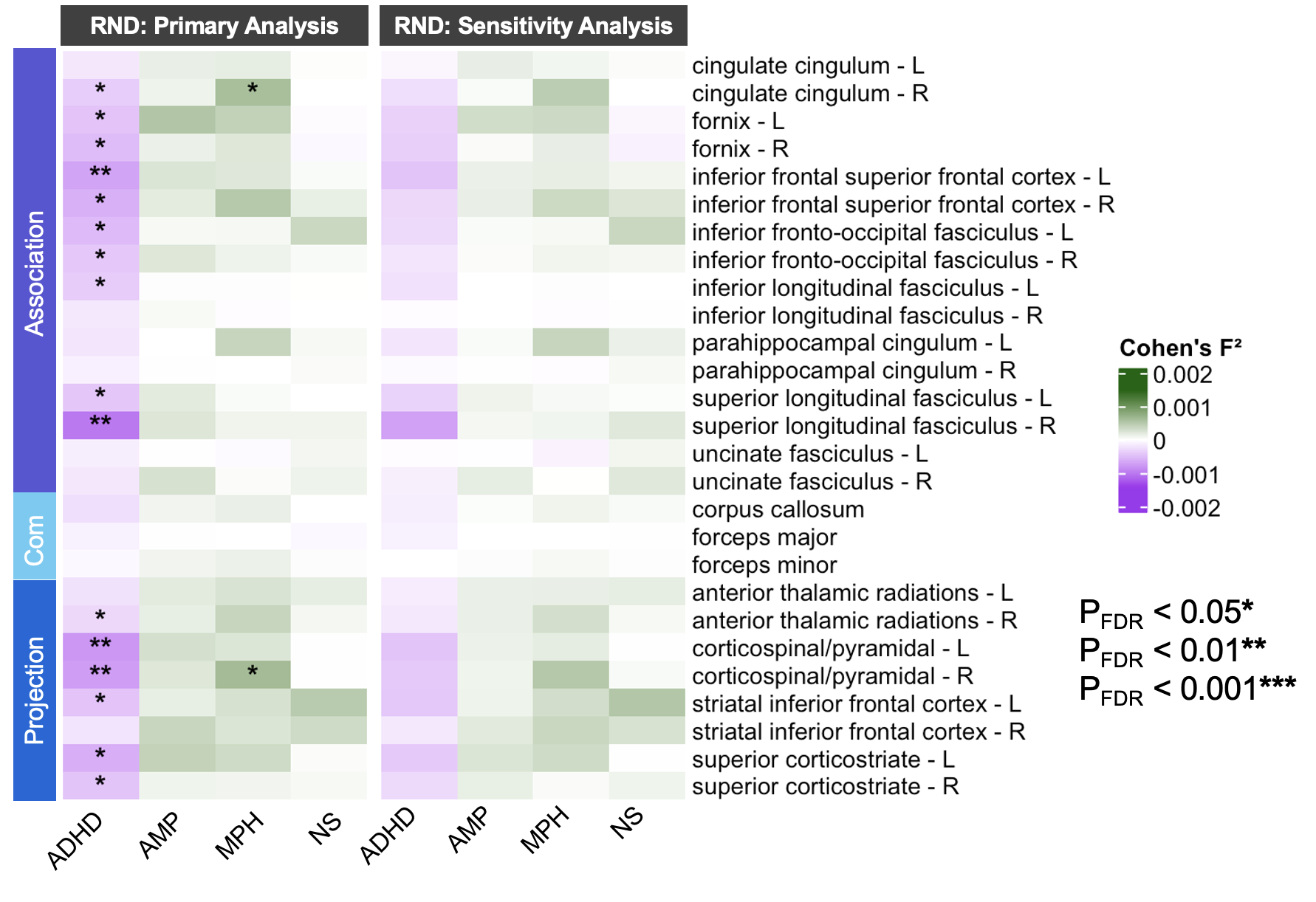
